# A novel anti-inflammatory treatment for bradykinin-induced sore throat or pharyngitis

**DOI:** 10.1101/2020.11.06.370395

**Authors:** Victor Leyva-Grado, Pavel Pugach, Nazlie Latefi

## Abstract

**Background:** Often thought of as a minor health concern, sore throat or pharyngitis is an important public health issue. It is one of the most common symptoms of upper respiratory diseases including COVID-19 and is a leading cause of physician visits and antibiotic prescriptions. However, few over the counter medications are proven to heal sore throat inflammation.

**Methods:** Adenocarcinomic human alveolar basal epithelial cells (A549 cells) and three dimensional organotypic human respiratory tissues were used to study inflammation and various treatment effects on respiratory epithelia. The cells and tissues were studied both in the presence and absence of bradykinin, one of the first inflammatory mediators of pharyngitis. Inflammation was measured by analyzing levels of prostaglandin E2 (PGE2), interleukin 8 (IL-8), and leukotriene B4 (LTB4), transepithelial electrical resistance (TEER), and lactate dehydrogenase (LDH) release. Tissue morphology was analyzed by immunohistochemistry.

**Results:** In studying pharyngitis using organotypic human respiratory tissue stimulated with bradykinin, we saw an increase in prostaglandin E2 (PGE2) and interleukin-8 (IL-8) in response to bradykinin. Acetyl salicylic acid (ASA), a non-specific COX inhibitor, was able to mitigate a bradykinin-induced increase in PGE2 in our studies. However, ASA was inflammatory above its therapeutic window, increasing levels of PGE2 and IL-8 above those seen with bradykinin stimulation alone. We describe a novel, scientifically validated treatment for sore throat, that contains a low dose of aspirin and other anti-inflammatory ingredients.

**Conclusion:** This study elucidates the complex mechanisms involved in healing pharyngitis, an inflammatory condition of the upper respiratory epithelia. An ASA-based formula (Biovanta) mitigated bradykinin-induced inflammation more strongly than ASA alone in organotypic human respiratory tissues. Surprisingly, we found that many of the most common over the counter sore throat therapies exacerbate inflammation and IL-8 in organotypic human respiratory tissues, suggesting these common treatments may increase the likelihood of further respiratory complications.

**Competing interest statement:** This study was funded entirely by Applied Biological Laboratories, a private company that owns the Biovanta™ product. Some studies were conducted by third parties in a blind format, as indicated. All other experiments were performed at Applied Biological Laboratories’ research facility located at the SUNY Downstate Biotechnology Incubator, a part of StartUP NY. All of the authors were employees of Applied Biological Laboratories at the time the experiments were performed.

## Background

Prior to the COVID-19 pandemic, pharyngitis, or sore throat was among the leading causes of physician visits and antibiotic prescriptions in the United States. This is surprising since viruses such as rhinovirus and influenza usually cause about 85% of sore throat cases (1). No over the counter cough and cold medicines have proven effective for sore throat either. Only systemically administered, non-steroidal anti-inflammatory drugs such as ibuprophen and paracetamol have proven effective for sore throat pain relief (2), but pharyngitis as the name implies also involves inflammation of the pharynx. Apart from gross physiological symptoms such as size of lymph nodes and throat color, it is very difficult to measure pharyngial inflammation in a human subject (3). Lack of reliable animal models may have also hampered drug discovery for this indication.

Fortunately, three-dimensional, organotypic tissue explants of human upper respiratory mucosa are now available (4-6). These models more reliably mimic the function of living tissue than either animal models or cell culture monolayers and can greatly accelerate drug discovery (4, 6). Using these models, we were thus able to study the biochemical pathway of pharyngitis in great detail and design a scientifically based treatment.

Bradykinin carried from the nasal passages to the nasopharynx during a respiratory infection can stimulate nociceptive receptors in the oropharynx and throat causing a sore throat (7-9). Bradykinin stimulation also increases IL-8, an inflammatory chemokine, and neutrophil chemoattractant often implicated as a component of “cytokine storms” in a time and concentration dependent manner in airway smooth muscle cells and respiratory epithelial cells via COX enzymes and PGE2 (10, 11). Clinically, at least three studies used bradykinin to induce sore throat in healthy subjects (8, 9, 12).

According to our findings, bradykinin does indeed increase inflammation in respiratory epithelia and organotypic cultures via the COX pathway. After an extensive literature search for compounds to mitigate this inflammation, we focused on acetyl salicyclic acid (ASA), or aspirin, a well-known, non-specific COX inhibitor. Aspirin is currently approved as an over-the-counter drug administered systemically via the oral route and as a general analgesic in the following dosages for adults: 325-650 mg every 4 hours, 325-500 mg every 3 hours, or 650-1000 mg every 8 hours.

We found that in A549 cells and in organotypic cultures, when applied apically, much lower dosages of ASA were needed, even after correcting for the *in vitro* testing conditions. There was a small therapeutic window in which ASA inhibited PGE2 release. Above this window, even in the absence of bradykinin, PGE-2 and IL-8 increased to significantly higher levels than baseline controls, creating an inflammatory state where we observed negative cytopathic effect.

When formulated with other anti-inflammatory compounds such as lysozyme, lactoferrin, and aloe, the anti-inflammatory effect of ASA at the therapeutic dose was increased, and greatly surpassed that of other leading sore throat remedies, which paradoxically, were found to be highly inflammatory and deleterious to the delicate respiratory lining. Organotypic cultures are a valuable research tool for this indication since the cytoarchitecture of the tissue mimics those in people, and we believe, effective dosages are likely to translate to clinically effective dosages.

## Methods

### Cells

A-549 cells (ATCC CCL-185) were cultured in DMEM Medium (Gibco) supplemented with 10% heat-inactivated fetal bovine serum (Gibco), 100 U ml−1 penicillin and 100 U ml−1 streptomycin (Gibco). Cells were seeded on 96 well cell culture plates at a density of 6400 cells/well. Before the experiment, when cells reached 90% confluency, media was replaced with serum free media. Human airway epithelia reconstituted in vitro were purchased from two different providers, EpiAirway™ tissues (AIR-100) from MatTek Corporation (Ashland, MA, USA) and MucilAir™ tissues from Epithelix Sàrl (Geneva, Switzerland). Specific cell culture media for each tissue were obtained from the respective manufacturer. Basal media was replaced with fresh media every other day until the day of the experiment.

### Reagents for treatment

ASA formula and ASA lozenge (Biovanta™ liquid and lozenge formulas respectively) are composed of the following ingredients: lactoferrin, lysozyme, acetyl salicylic acid, menthol, aloe and glycerin and can be purchased from leading pharmacies. Other over the counter treatments for sore throat and cold were obtained from a local pharmacy. Over the counter products in a lozenge presentation were prepared for use by dissolving the product in saliva buffer (13) in a weight/volume (w/v) ratio determined by the weight of the lozenge and the amount of water needed to dissolve that amount of the particular sugar in the lozenge. The weight volume ratios used are listed in Table 1. Products in liquid form were applied neat, except for the ASA formula because, unlike the other products, its indicated dosage is 10 times less than the average amount of saliva expected to be in the mouth. Recommended dosages are also listed in Table 1 and the salivary flow rate was assumed to be 1-2 ml/min. Three days before each study, the apical side of the EpiAirway™ and Mucilair™ tissue inserts was washed with 200 ul of media. Briefly, fresh-warmed media was gently added to the apical side of the insert to not disturb the tissues. The plates were incubated for 15 min at 37 C. After incubation, media from the apical side was gently pipetted up and down 3 times to remove the excess mucus formed on the tissue. The day of the experiment the inserts were transferred to new plates with fresh warmed media and then 10ul of each treatment was added to the cells or the apical surface of the human airway tissues, the plates were gently swirled and returned to 37C for 5 min. Every 24 hours, 10 ul of new treatment was applied. NS-398 (Sigma Aldrich) was applied apically as indicated in a volume of 10ul and at a concentration of 100uM diluted in saliva buffer. ASA (Sigma Aldrich) was applied apically at the indicated concentrations in a volume of 10ul at the indicated concentrations.

**Table 1.** Components and properties of tested sore throat treatments

| Product | Form | Active Ingredient | Dosage | Predominant Sugar | W/V Dissolution |
| --- | --- | --- | --- | --- | --- |
| **Theraflu Express Max Nighttime** | syrup | acetaminophen 650mg<br>diphenhydramine HCl<br>25mg phenylephrine<br>10mg | 30ml every 4 hours | n/a | n/a |
| **Mucinex Fast Max. DM Max** | syrup | dextromethorphan HBr<br>20mg, guaifenesin 400mg | 20 ml: 12mg-- every 4 hours | n/a | n/a |
| **Delsym** | syrup | dextromethorphan HBr<br>20mg, guaifenesin 400mg | 20 ml: 12mg-- no more than 6 doses in 24 hours | n/a | n/a |
| **Tylenol Cold and Mucus severe** | syrup | acetaminophen 325mg,<br>dextromethorphan HBr<br>10mg, guaifenesin 200mg,<br>phenylephrine HCl 5mg | 30ml every 4 hours | n/a | n/a |
| **Maty's cough syrup** | syrup | honey, apple cider vinegar, sea salt, clove, cinnamon, lemon balm, water, glycerin, lemon peel, marjoram, cayenne pepper | 1-2 teaspoons as needed | n/a | n/a |
| **Robitussin DM Max cough and chest congestion** | syrup | dextromethorphan HBr 20mg, guifenesin 400mg | 20ml every 4 hours | n/a | n/a |
| **Chloraseptic Max sore throat spray** | spray | phenol 1.5%, glycerin 33% | spit out after 15 seconds, use every 2 hours | n/a | n/a |
| **Biovanta throat spray** | spray | acetyl salicylic acid 6mg, menthol 5mg | apply one spray from each side to throat, repeat up to once every 30 min | n/a | n/a |
| **Ricola original** | lozenge | menthol 4.8mg | dissolve 2 drops in mouth, repeat every 2 hours as needed | starch syrup, sugar | 1:0.75 |
| **Ricola Dual Action sore throat and cough honey lemon** | lozenge | menthol 8.3 mg | dissolve 1 drop in mouth, repeat every 2 hours as needed | starch syrup, sugar | 1:0.75 |
| **Halls Relief honey lemon** | lozenge | menthol 7.5mg | dissolve 1 drop in the mouth every 2 hours | glucose syrup, sucralose, sucrose | 1:0.75 |
| **Vicks Vapacool severe** | lozenge | menthol 20mg | dissolve in mouth, repeat as necessary | corn syrup | 1:1 |
| **Ludens** | lozenge | pectin 2.8mg | dissolve in mouth, repeat as necessary | corn syrup, sucrose | 1:0.75 |
| **Coldeeze** | lozenge | zincum gluconicum 2X (13.3mg zinc) | dissolve in mouth, repeat every 2-4 hours as needed | corn syrup, honey, glycine, sucrose | 1:0.75 |
| **Cepacol Instamax** | lozenge | benzocaine 15mg, menthol 20mg | 1 lozenge every 2 hours | isomalt, maltitol, propylene glycol | 1:1.4 |
| **Strepsils sore throat and cough** | lozenge | dichlorobenzyl alcohol, amylmetacresol, and levomenthol | 1 lozenge every 3 hours | sucrose and glucose | 1:0.75 |
| **Wedderspoon Manuka Honey** | lozenge | honey | n/a | cane sugar (sucrose) | 1:0.75 |
| **Zarbees 96% Honey Cough Soothers natural lemon menthol** | lozenge | english ivy leaf extract 20mg | dissolve lozenge in mouth, repeat as needed | honey | 1:0.6 |
| **Zicam cherry** | lozenge | Zinc (10mg) | 1 lozenge every 3 hours as needed | corn syrup, sucralose | 1:1 |
| **Fisherman's Friend** | lozenge | menthol 10mg | 1 lozenge every 2 hours as needed | dextrin, licorice, sugar, tagacanth | 1:1 |
| **Biovanta lozenge** | lozenge | acetyl salicylic acid 6mg, menthol 5mg | 1 lozenge up to every 30 minutes as needed | isomalt | 1:1.4 |

### Bradykinin Challenge

Cell cultures and tissues were challenged with different concentrations of bradykinin from Tocris, Minneapolis, MN. Bradykinin was diluted to the appropriate concentration using saliva buffer (13). After tissues were treated, ten microliters of the bradykinin solution were added to the cells (A549 cells) or the apical surface of the human airway tissues. Plates were gently swirled and returned to 37C incubation. Samples from basal media were collected at different time points after challenge and stored at -80C until use. Bradykinin treatment was not repeated after the initial application.

### Measurement of Transepithelial Electrical Resistance (TEER)

TEER was measured in the human airway epithelium inserts to determine the integrity of tight junctions with a Millicell ERS-2 volt-ohmmeter (Millipore Sigma, Burlington, MA, USA). Briefly, inserts were transferred to new plates with 600 ul of warmed media per well, then 200 ul of media were gently added to the side wall of the inserts apical side. Plates were incubated for 5 min at 37C and then the TEER measurements were collected. Resistance values (Ω) were converted to TEER (Ω.cm2) by using the following formula: TEER (Ω.cm2) = (resistance value (Ω) -100(Ω)) x 0.33 (cm2), where 100□Ω is the resistance of the membrane and 0.33□cm2 is the total surface of the epithelium.

### Cytotoxicity

Cytotoxicity was assessed via lactate dehydrogenase (LDH) concentrations measured in 100 ul of basolateral medium incubated with the reaction mixture of a cytotoxicity detection kit (Sigma-Aldrich, Roche, Saint Louis, MO, USA) following the manufacturer’s instructions. To determine the percentage of cytotoxicity, the following equation was used (A = absorbance values): Cytotoxicity (%) = (A (exp value)-A (low control)/A (high control)-A (low control))*100. The high control value corresponds to a 10% Triton X-100 treatment applied to the culture for 24 h. A threshold limit of 5% of the toxicity index reflects the physiological cell turnover in human airway epithelium cultures.

### Analysis of inflammatory response

To evaluate the inflammatory response, cell culture supernatant and basolateral media from tissues was collected at different time points after challenge. Cyclooxygenase-2-derived prostaglandin E2 (PGE2) and LTB4 release were measured with enzyme-linked immunosorbent assay kits (ELISA; R&D systems, Minneapolis, MN) according to the manufacturer’s instructions. Interleukin-8 (IL-8 or CXCL8) protein in the basolateral media was measured using a magnetic bead-based ELISA (Procartaplex, ThermoFisher Scientific, Waltham, MA) according to the instructions provided by the manufacturer and the plate were then read using a Luminex-based Bio-plex Multiplex system (Bio-Rad, Hercules, CA). Samples were diluted to 1:3 for PEG− 2 and LTB4 analysis and 1:20 for IL-8 measurements.

### Histopathology

At the end of the experiment, inserts with the human airway epithelia cells were fixed with a formalin solution (neutral buffered, 10%) and kept at 4C until being sent to Histowiz (New York, NY, USA) for histology processing. Inserts were bisected to allow histological staining of paraffin-embedded sections. Paraffin-embedded tissues were cut into 5-µm sections, de-paraffinized, and the rehydrated sections were stained with hematoxylin–eosin. The images were collected from digitalized slides at a final magnification of 20X.

### Statistical analysis

Data are presented as mean ± standard error of the mean. For statistical comparison of differences between groups, results were analyzed by an unpaired t test, Mann-Whitney, ANOVA or Kruskal-Wallis tests using the GraphPad Prism software (version 6.01, La Jolla, USA). A p-value ≤ 0.05 was considered significant.

## Data availability

All the presented data is accessible on the Open Science Framework https://osf.io/ejcf8/

## Results

Bradykinin, one of the first chemokine signals for pharyngitis causes inflammation in A549 cells that can be blocked by acetyl salicylic acid (ASA) Bradykinin increases PGE2 via the arachidonic acid-COX 2 pathway, and acetyl salicylic acid (ASA) is known to be a potent COX inhibitor (11). First, we sought to determine the effects of bradykinin on <u>adenocarcinomic human alveolar basal epithelial cells (</u>A549 cells). Various concentrations of bradykinin were applied to A549 cells to assess the downstream effects on PGE2 and IL-8 (Figures 1A and 1B). LDH was also measured to assess cytotoxicity (Figure 1C).

**Figure 1.**
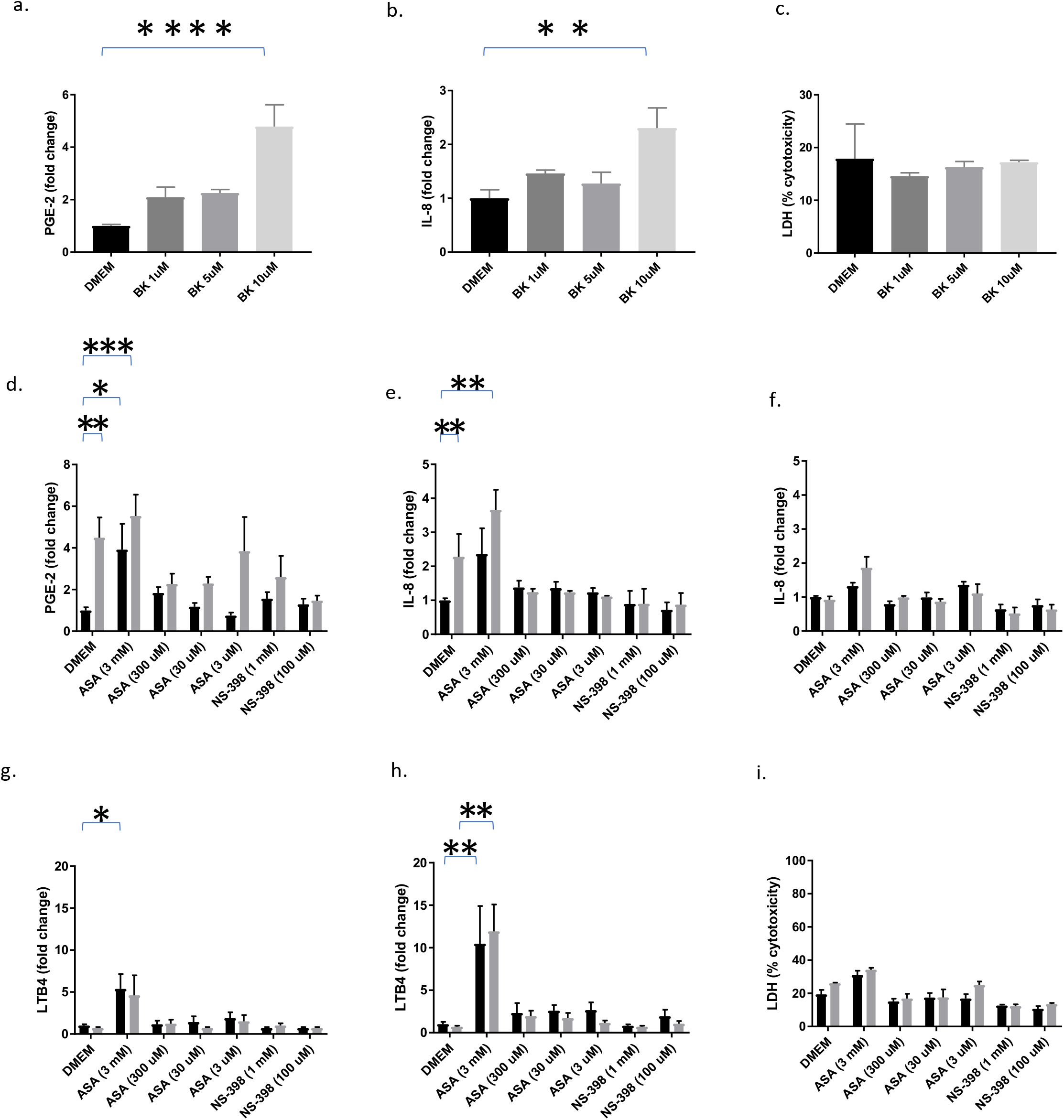
A-C: A549 cells were treated with culture media (DMEM), or the indicated concentrations of bradykinin (BK) in 96-well microtiter plates. PGE2 levels in media were measured at 4 hours post-inoculation. IL-8 was measured 24 hours post inoculation (B). LDH levels were measured 24 hours post-inoculation (C). D-I: A549 cells were treated with cell culture medium (black bars) or 10uM of BK (gray bars) 30 minutes after treatment with various concentrations of ASA or 100uM NS-398. PGE2 was measured at 4 hours post-inoculation (D). IL-8 was measured 24 (E) or 48 (F) hours post-inoculation. LTB4 was measured 4 (G) and 24 (H) hours post-inoculation. LDH was measured 48 hours post-inoculation (I). Mann-Whitney statistical tests were performed using GraphPad Prism software.

At least 10uM bradykinin (supplied by Tocris) was needed to stimulate a statistically significant increase in PGE2 compared to negative control, about a 5-fold increase (Figure 1A). This dose of bradykinin has been shown by others to elicit significant increases in PGE2 and Il-8 in airway smooth muscle cells (10) and A549 cells at 4 hours and 24 hours respectively (11). As a result of bradykinin stimulation, there was also a statistically significant increase in IL-8 expression after 24 hours (Figure 1B). There was no observable cytopathic effect from the increase in PGE-2 and Il-8 expression, and no discernable change in LDH release was observed (Figure 1C).

Once we established the stimulatory effects of bradykinin, we asked whether it could be inhibited by acetyl salicylic acid (ASA), a non-specific COX-inhibitor, or NS-398, a COX-2 specific inhibitor. Prior to stimulating with 10uM bradykinin, we treated cells with various concentrations of ASA, or NS-398 (Figure 1D-I grey bars). For cells treated with ASA: the lowest concentration, 0.6 ug/ml (3 uM) did not affect the bradykinin-induced increase in PGE2 secretion, but it did bring IL-8 down to baseline (Figure 1 D, E, and F). The medium concentrations, 6 ug/ml (30 uM) and 60 ug/ml (300 uM) of ASA mitigated the effects of bradykinin on PGE2 and IL-8, bringing both down to baseline (Figure 1D, E, and F). The highest 600 ug/ml (3mM) dose of ASA was inflammatory, leading to an increase of PGE2 and IL-8 over levels reached with bradykinin stimulation alone (Figure 1D, E, and F).

To be sure the inflammatory effect of high ASA was not due to an interaction with bradykinin, we also measured levels of PGE2 and IL-8 in response to various concentrations of ASA and NS-398 in the absence of bradykinin (1D-I black bars). The highest level of ASA tested, 3 mM, was indeed inflammatory even in the absence of bradykinin. It brought levels of PGE2 up 4-fold compared to baseline, and levels of IL-8 up 2-fold over baseline (Figure 1D and E). It also increased secretion of LDH, a marker of toxicity (Figure 1I). The lower concentrations of ASA were not inflammatory on their own, and neither were any of the concentrations of NS-398 tested.

To help elucidate the reason for the toxicity of higher levels of ASA, we measured levels of LTB4 in response to the various concentrations of ASA and NS-398 in the presence and absence of bradykinin (1G-I). While bradykinin alone did not stimulate LTB4 release, 3 mM ASA did. Possible explanations for this will be described in the Discussion section.

The bradykinin-induced inflammatory cascade can be replicated in an organotypic model of the human respiratory system MucilAir™ is composed of basal cells, ciliated cells and goblet cells that secrete mucus. The cells used to prepare these organotypic cultures are derived from nasal polyp biopsies. The proportion of the various cell types is preserved compared to what one observes *in vivo* (14) and the cytoarchitecture is virtually identical to that of the pharynx (15). We chose to use the organotypic tissue model because we hypothesized that they would be more predictive of human efficacy than animal models (4,6). These tissue models secrete most of the same cytokines that are secreted *in vivo*, and since the treatments we are designing are intended to be used topically, the metabolism and targeting of the ingredients should be virtually identical to what would be observed *in vivo* (6, 16).

We determined 47mM (50mg/ml) to be the lowest effective dose of bradykinin that would reliably elicit a significant response of PGE2 in Mucilair™. Next, we investigated whether various concentrations of ASA could inhibit bradykinin-induced inflammation in Mucilair™. In vivo, ASA would be taken by mouth and then diluted by saliva before reaching the pharynx, where bradykinin is released physiologically. Therefore, after assuming a 1000-fold increase from A549 cells, where we found 300uM and 30uM to be most effective, we assumed a 10-30-fold dilution for salivary flow (for a formula dosed every 10-30 min and a salivary flow rate of 1-3 ml/min), which would ultimately give us a 100-fold increase in concentration from A549 cells. Hence, we tested 3 mM (0.6 mg/ml) and 33 mM (6 mg/ml) of ASA with 47 mM of bradykinin in the MucilAir™ organotypic model. We measured levels of PGE2, and IL-8, both in the presence and absence of bradykinin (Fig 2 A-E); black bars (no bradykinin), and grey bars (with bradykinin).

**Figure 2.**
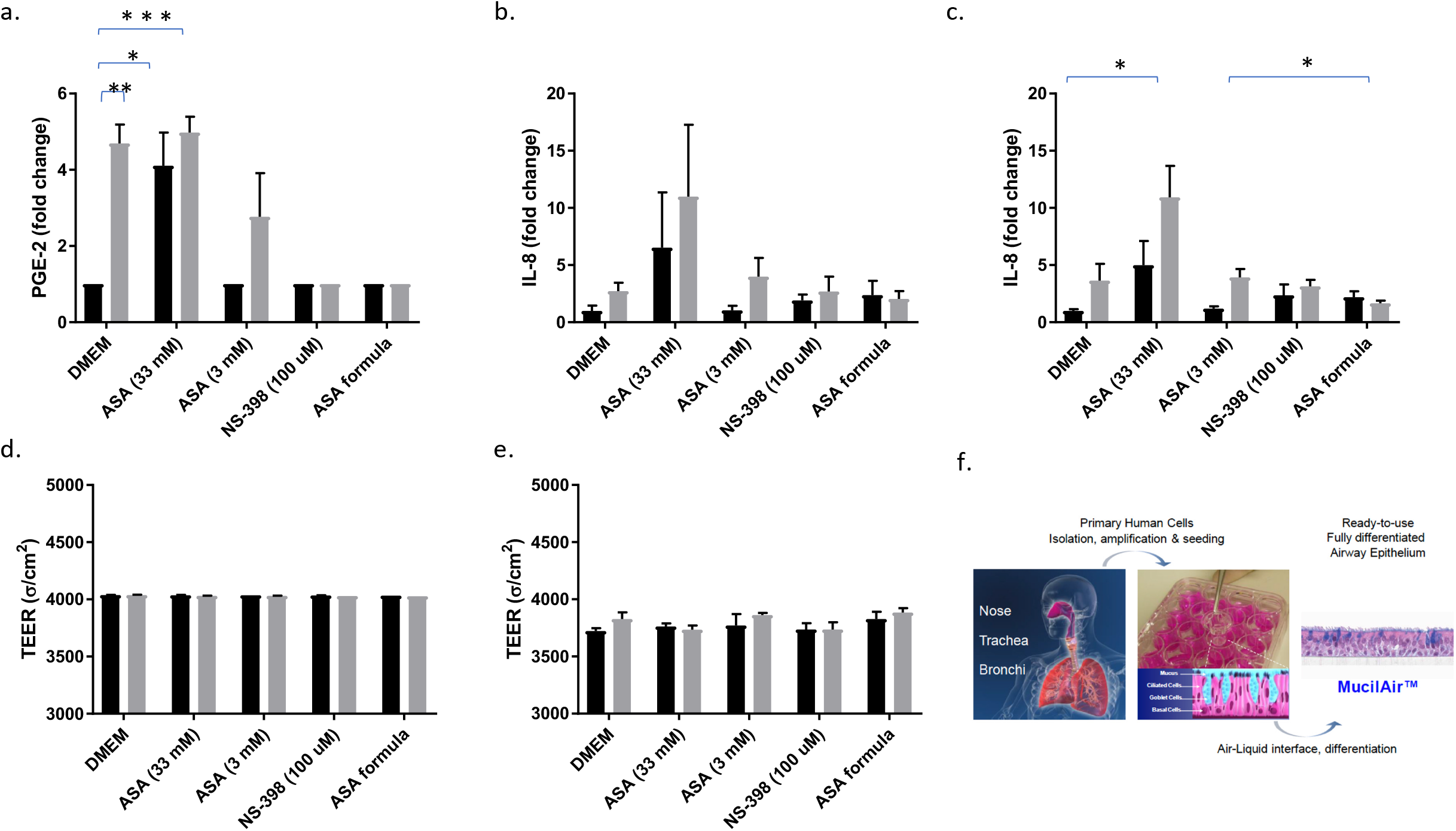
Mucilair™ respiratory tissues suspended in transwell inserts were treated with either saliva buffer (vehicle, black bars), or 47 mM bradykinin (gray bars) following treatment with various concentrations of ASA, 100mM NS-398, or an ASA-based formula containing 3 mM ASA (Biovanta liquid). Levels of PGE2 in basal media were measured at 4 hours post-inoculation (A). IL-8 levels in basal media were measured at 24 (B) and 48 (C) hours post-inoculation. Transepithelial membrane resistance was measured at 24 (D) and 48 (E) hours post-inoculation (C). Statistics shown are results of unpaired t-test using GraphPad Prism software. Statistics were performed on raw data without omissions and not on fold-change values. The figures in D are reprinted with permission from Epithelix Sarl.

Bradykinin stimulation led to a statistically significant increase in PGE2 at 4 hours, as did stimulation with the higher, 33 mM concentration of ASA both in the presence and absence of bradykinin (Fig 2A), confirming our findings from A549 cells that higher concentrations of ASA can be inflammatory. The lower 3 mM concentration of ASA was not inflammatory on its own, but it did prevent levels of PGE2 from rising to levels reached with bradykinin stimulation alone (Fig 2A). Neither NS-398, nor the ASA formula (Biovanta liquid containing 3mM ASA) increased PGE2 on their own, however both did successfully keep PGE2 levels from rising in the presence of bradykinin (Fig 2A). At 24 hours, the higher, 33 mM concentration of ASA increased IL-8 levels both on its own and in the presence of bradykinin. The effects were visible but did not reach statistical significance (Fig 2B). At 48 hours, stimulation with 33 mM ASA increased IL-8 but it only reached statistical significance in the presence of bradykinin. Neither NS-398, nor the lower 3 mM concentration of ASA increased IL-8 above control levels. Transepithelial electrical resistance (TEER) measures at 24 and 48 hours did not indicate membrane damage with any of the treatments (Fig 2 D, E).

### A Novel, ASA-based formula is more effective than ASA alone at blocking bradykinin-induced PGE2 production in epithelial tissues

Next, we sought to investigate the anti-inflammatory effects of an ASA-based formula (Biovanta™ throat spray), a formula containing 30 mM ASA (which is diluted 10 fold upon treatment), lysozyme, lactoferrin, aloe, glycerin, and menthol following bradykinin stimulation. Lysozyme and lactoferrin are potent anti-inflammatory molecules present in human nasal secretions. They are also used as preservatives and excipients to affect the rheological properties and viscosity of liquid formulations. Aloe, glycerin, and menthol are common excipients used in cough and cold products, are all natural, and were determined to not be toxic at the concentrations used (data not shown). We hypothesized that they could have beneficial anti-inflammatory effects.

We pretreated the tissues with either buffer, ASA alone, the ASA-based formula (Biovanta throat spray), or NS 398, prior to treating with bradykinin. Indeed, at 48 hours after bradykinin stimulation, tissues treated with Biovanta showed a statistically significant decrease in IL-8 levels compared to those treated with only 3 mM ASA. Tissues treated with Biovanta also seemed to show a bigger decrease in PGE2 4 hours after stimulation than those treated with 3 mM ASA alone, however the difference was not statistically significant.

### An ASA-based formula (Biovanta throat spray) containing anti-inflammatory excipients decreases PGE2 in Mucilair™ without the cytotoxicity, or membrane damage caused by leading sore throat products

After establishing the superiority of the ASA-based formula against ASA, we compared its efficacy in decreasing bradykinin-induced inflammation to leading sore throat products. Bradykinin at a concentration of 47 mM was used to induce inflammation after pretreating the tissues for five minutes with either buffer, 100 uM NS-398, an ASA-based formula (Biovanta™ throat spray), or various over the counter sore throat products containing the following active ingredients: phenol and glycerin (Chloraseptic max sore throat spray), dextromethorphan and guifenesin (Robitussin DM max), english ivy leaf extract (Zarbees), or menthol (Halls relief honey lemon). In tissues pre-treated with buffer, PGE2 increased 1.54-fold after 4 hours and IL-8 increased 1.75-fold after 24 hours and remained high at 48 hours (Figure 3A-C). All three increases were statistically significant compared to negative control (p=0.02 and p=0.04, respectively). Pretreatment with either NS-398 or the ASA formula kept PGE2 at control levels, while three of the other sore throat treatments increased PGE2 levels, albeit not to statistically significant levels (Figure 3A). At 24 hours post-stimulation, bradykinin stimulation increased IL-8 levels significantly (p=0.0352). IL-8 levels in tissues pretreated with either NS-398 or the ASA formula did not increase, they were similar to tissues pretreated with vehicle.

**Figure 3.**
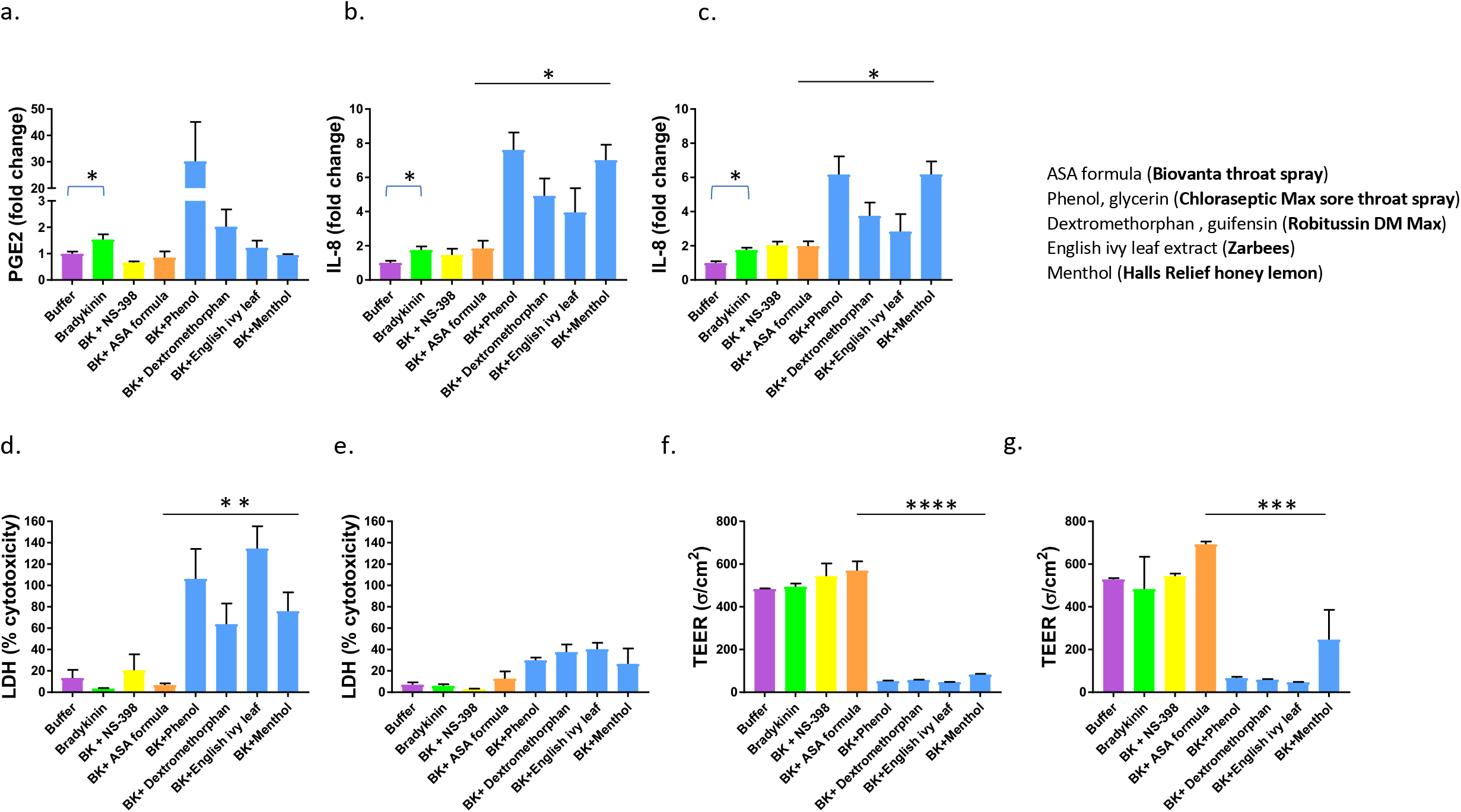
Mucilair™ respiratory tissues suspended in transwell inserts in 24 well plates were treated with saliva buffer (vehicle), bradykinin (BK) (47 mM) or BK following treatments with 100uM NS-398 or various over the counter sore throat products. The products are indicated on the graph by their first active ingredient. The key lists all the active ingredients and the trade name of the product. Levels of PGE2 in basal media were measured at 4 hours post-inoculation (A). IL-8 levels and LDH in basal media were measured at 24 hours (B and D) and 48 hours (C and E) post-inoculation, and transepithelial electrical resistance was measured at 24 hours (F) and 48 hours (G) post inoculation. Statistics shown are results of ANOVA and unpaired t-tests using GraphPad Prism software. Statistical analysis was performed on raw data without omissions and not on fold-change values.

Of the leading sore throat products tested in this experiment, three were liquid formulations (either sprays or syrups) and one was a popular lozenge. None of the products prevented an increase in either PGE2 or IL-8. In fact, paradoxically, phenol and glycerin (Choloroseptic® Max Sore Throat Spray) and dextromethorphan and guifenesin (Robitussin®DM Max) pretreatment increased PGE2 levels compared to buffer pretreatment (Figure 3A). In addition, all of the products caused significant cytotoxicity as measured by increased IL-8 secretion, increased LDH secretion, and lowered TEER (Figure 3B-G). All four products tested showed statistically significant increases in IL-8 compared to ASA formula (Biovanta throat spray) (p=0.0142 after 24 hours and p=0.0140 after 48 hours), statistically significant increases in LDH cytotoxicity (p=0.0094 after 24 hours), and statistically significant decreases in TEER (p<0.0001 after 24 hours and p=0.0001 after 48 hours).

### The inflammatory effects of leading over the counter sore throat products were confirmed in the EpiAirway™ model in the absence of bradykinin, while neither an ASA-based formula nor a lozenge was inflammatory

The following products were applied to the apical surface of EpiAirway™ to assess inflammatory effects: an 8.3mg menthol containing lozenge (Ricola® dual-action sore throat and cough honey lemon), a pectin containing lozenge (Luden’s® wild cherry flavor dual action), a 13.3mg zinc containing lozenge (Cold-EEZE natural cherry flavor lozenge), a honey containing lozenge (Wedderspoon organic manuka honey drops), a 10mg zinc containing lozenge (Zicam® cherry lozenge), a dichlorobenzyl alcohol, amylmetacresol, and levomenthol containing lozenge (Strepsils® sore throat and cough lozenges), a 20mg menthol containing lozenge (Vicks® Vapacool severe drops), an ASA-based formula (Biovanta throat spray), and an ASA-based lozenge (Biovanta lozenge). These products were applied to tissues after being dissolved in saliva buffer. The lozenges were weighed and dissolved in a predetermined weight/volume (w/v) ratio of saliva buffer. Since hard lozenges are amorphous solids and about 98-99% sugar (17) the w/v ratio needed to dissolve each lozenge was calculated as the minimum volume needed to dissolve the predominant sugar in the lozenge based on its solubility and density. The dilution volume for each lozenge tested is listed in Table 1.

The ASA-based formula (Biovanta™ throat spray) is an over-the-counter drug and is intended to be sprayed to the back of the throat in a total volume of 200 ul. We considered the average salivary flow rate to be 2 ml/min and that the liquid formula would be diluted 1:10 by saliva in the mouth. Therefore, we diluted the liquid formula 1:10 before applying it to the tissues. The other liquid spray formulas and syrups tested had dosage volumes far exceeding the amount of saliva likely to be present in the mouth at any given time, so they were applied neat. After applying 10ul of the dissolved lozenge or ASA-based formula (Biovanta throat spray) to the tissues, the tissues were placed at 37 degrees Celsius for 5 minutes and then 10ul of saliva buffer was applied to the tissues.

Treatments were reapplied every 24 hours. The tissues were not stimulated with bradykinin. As illustrated in figure 4, except for the ASA-based formula and lozenge (Biovanta throat spray and Biovanta lozenge), all the products tested caused increases in inflammatory cytokines (PGE2, IL-8, or both), and LDH and also disrupted membrane integrity as illustrated by significantly lower TEER measurements. Note that although some products appear to decrease IL-8 levels at 24 hours, the same products show significant decreases in TEER and increases in LDH, suggesting that low IL-8 levels may be due to cell death. All nine products tested showed statistically significant changes in PGE2 levels compared to ASA formula (Biovanta throat spray) and ASA lozenge (Biovanta lozenge (p=0.0013), statistically significant changes in IL-8 levels (p=0.0010), statistically significant changes in LDH levels (p=0.0033), and statistically significant decreases in TEER (p=0.0015).

**Figure 4.**
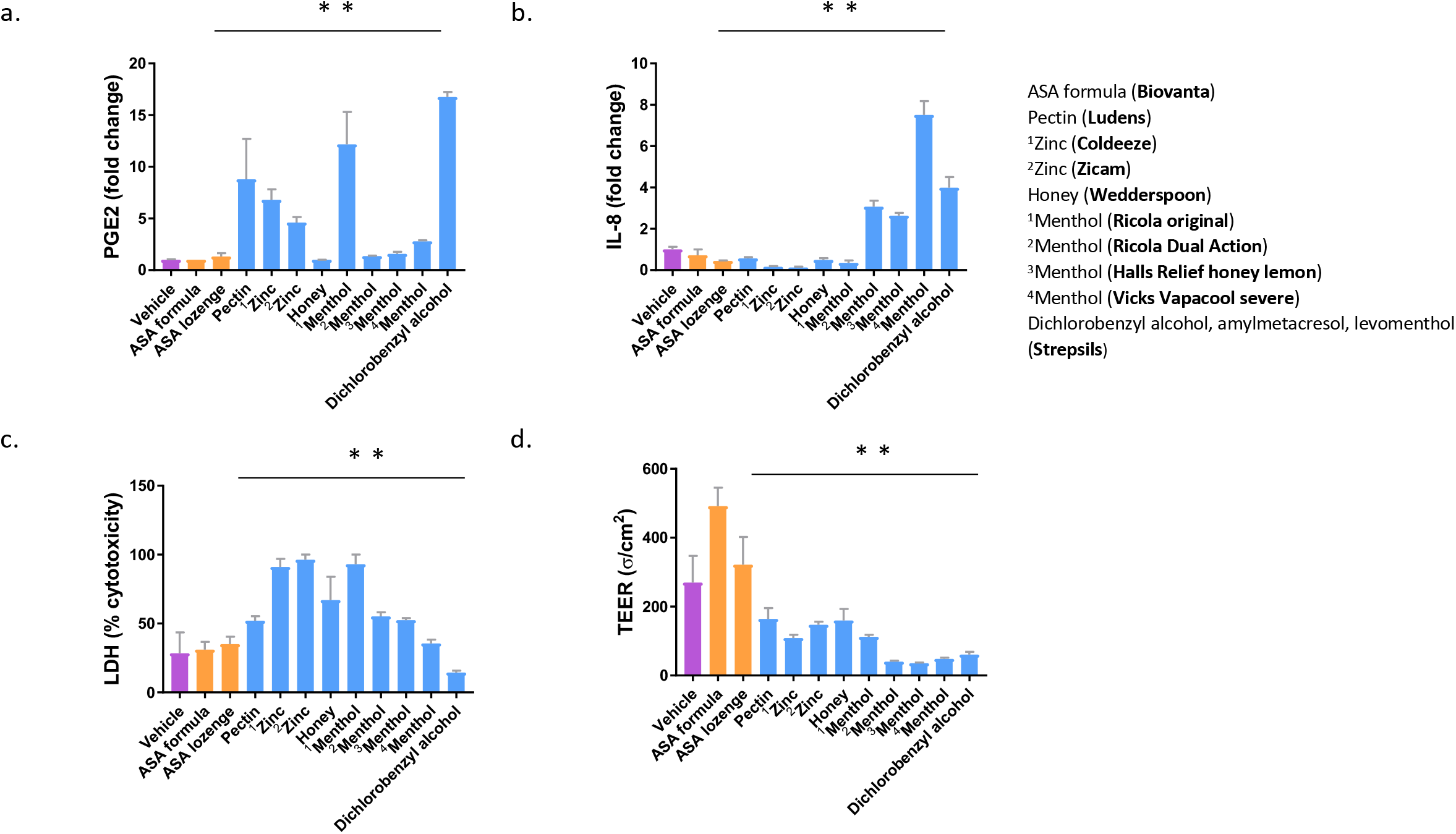
EpiAirway™ (AIR-200-PE6.5) respiratory tissues suspended in transwell inserts were treated with saliva buffer (vehicle) and various sore throat treatments as indicated. The products are indicated on the graph by their first active ingredient. The key lists all the active ingredients and the trade name of the product. Measurements of PGE2 at 4 hours (A), IL-8 at 24 hours (B), LDH at 24 hours, (C) and TEER at 24 hours (D) are shown. Statistics shown are results of Kruskal-Wallis tests performed using GraphPad Prism software. Statistics were performed on raw data without omissions and not on fold-change values.

Neither the ASA-based formula (Biovanta throat spray), nor the ASA-based lozenge (Biovanta lozenge) were inflammatory. In fact, levels of PGE2, IL-8, LDH, and TEER in response to Biovanta were comparable to the vehicle condition (Figure 4 A-D).

### According to a third-party blind placebo-controlled study, leading sore throat and cold remedies physically damage respiratory epithelia, but the ASA lozenge (Biovanta lozenge) does not

Unlabeled Eppendorf tubes containing the various sore throat syrups and lozenges were sent to Mattek Corp (Ashland MA) for third party blind analysis. EpiAirway™ (AIR-200-PE6.5) were treated in triplicate with each of the syrups (undissolved) and each of the lozenges dissolved in buffer. Following a 5 min incubation, a 15 mM solution of bradykinin was added to the inserts. We found that in these tissues 15 mM bradykinin was sufficient to increase PGE2 levels (data not shown). After incubating at 37C for 48 hours, TEER measurements were taken. Of the products tested, only the isomalt-based, ASA lozenge (Biovanta lozenge) did not induce measurable damage to epithelial tissues (Figure 5).

**Figure 5.**
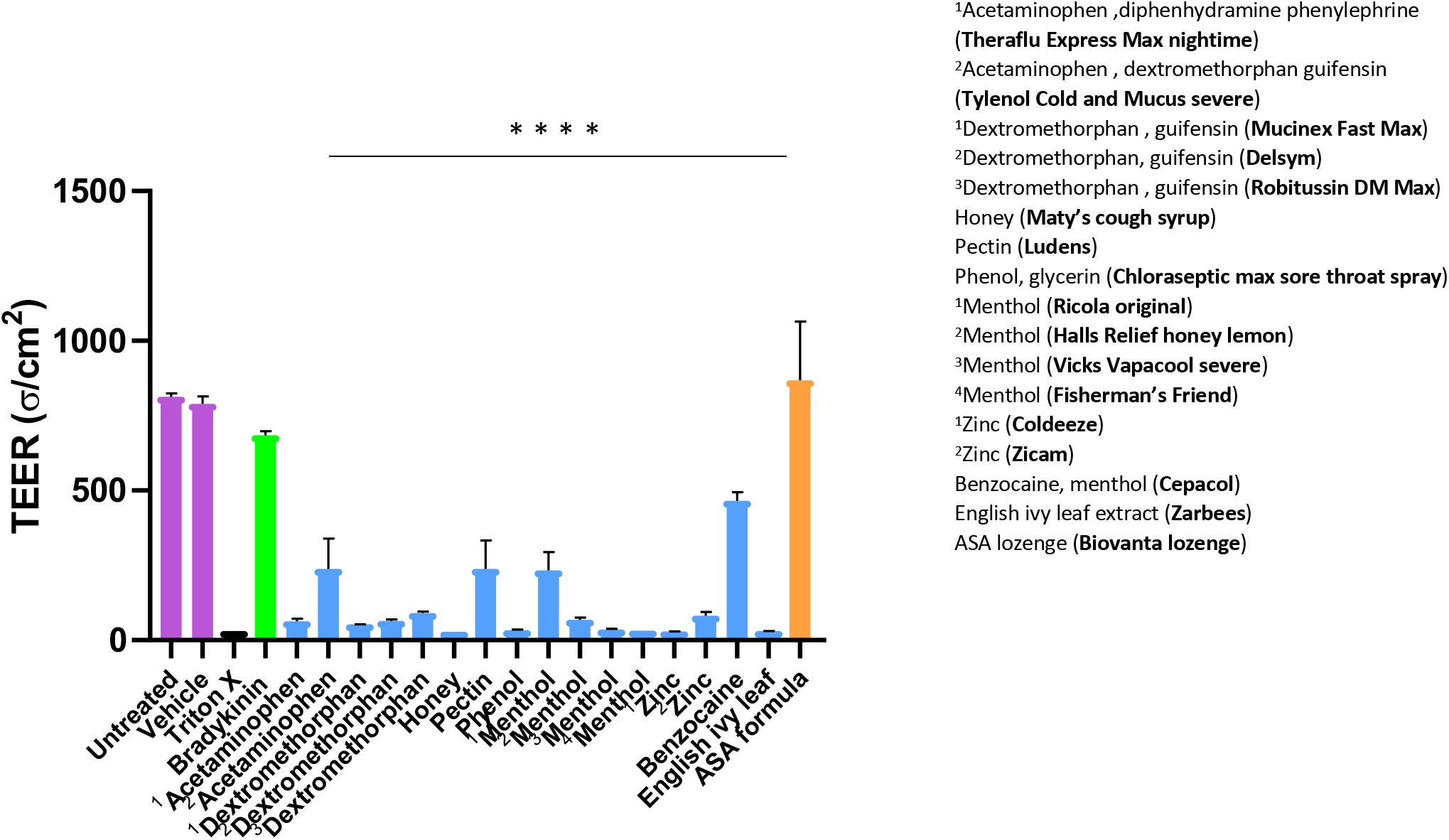
This study was performed in a third party blind format. EpiAirway™ (AIR-200-PE6.5) respiratory tissues suspended in transwell inserts were either left untreated, treated with saliva buffer (vehicle), triton-X (positive control), or various sore throat products as indicated. The various products are indicated on the graph by their first active ingredient. The key lists all the active ingredients and the trade name of the product. Following their respective treatments all tissue inserts (except the untreated) were treated with 15mM bradykinin. TEER was measured 48 hours after treatment. Statistics shown are results of a Kruskal-Wallis test using GraphPad Prism software, p<0.0001. Statistics were performed on raw data without omissions and not on fold-change values.

Considering an average salivary flow rate of 2 ml/min and an average recommended dosage for throat syrups of 20-30 ml, we don’t believe the syrups are intended to be dissolved in saliva but instead to bathe the oral pharynx. The lozenges were dissolved in buffer at a concentration determined by their weight and the type(s) of sugars they contain. For example, Halls® Relief Honey Lemon contains mainly glucose, according to its label. Glucose has a solubility of 91g/100ml of solution and a density of 1.02 g/ml. Each lozenge weighs 3.15 g. It would therefore take about 1:0.75 w/v of lozenge to buffer (or saliva) to dissolve each lozenge. See table 1 for a list of the various products and the w/v ratio of buffer they were dissolved in.

### Histological analysis confirms extensive tissue damage by leading products, as measured by TEER

Following the 48-hour experiment depicted in Fig 5, the organotypic respiratory tissues were fixed in 10% paraformaldehyde, embedded in parrafin, and sectioned into 5 um thick sections taken from the center of each section after it was cut in half. The sections were then treated for hemotoxylin and eosin (H&E) staining to observe any histological changes. The average thickness of the untreated sections was 55-60 um from the top of the cilia to the bottom of the basal cell layer. The groups treated with the most common over the counter sore throat products exhibited marked loss of cilia and pseudostratified columnar epithelial cells (Figure 6). In most cases, the columnar epithelial cells showed atrophy which decreased the thickness of the epithelia to about half of control values.

**Figure 6.**
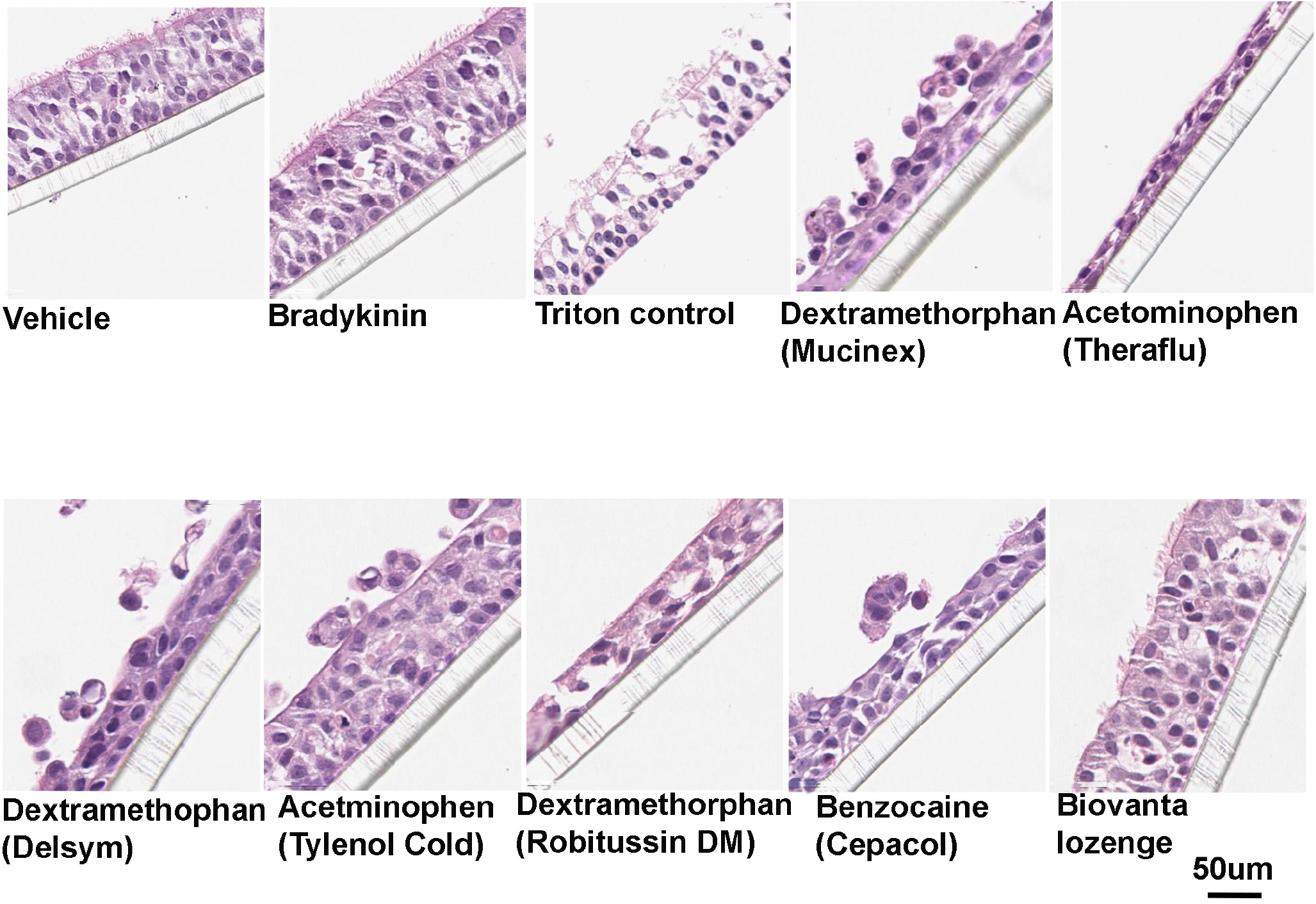
Histological analysis of the tissues analyzed in the third-party blind study, a representative sample of the tissues used in figure 5 are shown. In all of the tissues, except those treated with saliva buffer (vehicle), bradykinin, or the ASA lozenge (Biovanta lozenge), marked histological damage was seen in response to treatment. After fixation and embedding in parrafin, sections were cut into 5 um thick section and stained with hemotoxylin and eosin.

## Discussion

Despite the plethora of over the counter (OTC) products and supplements available to the US consumer, none have shown efficacy for sore throat in clinical studies. A Cochrane meta-analysis found that non-steroidal anti-inflammatory drugs (NSAIDS) such as aspirin when taken orally are “somewhat effective” in relieving the discomfort caused by a cold (18). Considering the mechanism by which bradykinin induces inflammation through the COX-1/COX-2 pathway, it is surprising that systemically administered NSAIDs have not proven to be more effective. Perhaps, they need to be applied locally or at different dosages. Indeed, we found that there is a small therapeutic window at which aspirin decreases inflammation and above which it becomes inflammatory.

At the therapeutic dose, we found that ASA and an ASA formula (Biovanta throat spray) blocked the bradykinin-induced increase in PGE2. We expected to also see a decrease in IL-8 but this was only observed in A549 cells, and not in Mucilair. IL-8 levels did not reach statistical significance above baseline in Mucilair, and there was therefore not much room for significant decreases. Mucilair, an organotypic model, contains several cell types and numerous cytokines that might modify the increases in IL-8 expression seen in A549 cells.

At a dose 10-fold higher than the effective dose, aspirin caused a statistically significant 4-fold increase in PGE2 above baseline in A549 cells and in Mucilair, as well as a statistically significant 2-fold increase in IL-8 above baseline in A549 cells, and a 5-fold increase above baseline in Mucilair which did not reach statistical significance. A dose-dependent effect of aspirin has been detected in several different physiological systems (19). Our results are consistent with a clinical study showing that a moderate dose (500 mg/day) of systemically administered aspirin, which is much lower than the recommended dose for adults, effectively treated ACE-inhibitor induced cough, which is thought to be mediated by bradykinin (20).

In order to shed some light on what might be driving the increase in PGE2 and IL-8 expression with high doses of ASA, we measured levels of LTB4 in response to high dose ASA in the presence and absence of bradykinin. High dose, 3 mM ASA caused a statistically significant, 5-fold increase in LTB4 above baseline both in the presence and absence of bradykinin in A459 cells.

Apart from the COX pathway, arachidonic acid is also responsible for the generation of leukotrienes via lipoxygenase. Inhibiting the COX pathway with COX inhibitors may shunt arachidonic acid to the highly inflammatory lipoxygenase pathway (14, 21), a pathway, which coincidentally, may be responsible for bronchoconstriction in aspirin induced asthma (21, 22). We observed an increase in one of the major lipoxygenase products, LTB4, as well as IL-8 and PGE2 in A549 cells stimulated with high concentration ASA. Prior studies confirm that LTB4 stimulates IL-8 (10, 23). LTB4 also activates several kinase cascades leading to production of several cytokines, many of which might increase PGE2 (24). For example, TNFα and IL-1β were shown to increase PGE2 expression in A549 cells (25).

Interestingly, the highest dose of NS-398 tested did not show any inflammatory effect. This may be due to its selectivity for COX-2, therefore leaving some COX-1 available to produce prostaglandins. Alternatively, since A549 cells greatly over-express COX-2, it may not be possible to knock down levels completely. Other groups have shown that A549 cells express both COX-1 and COX-2 mRNA and that over-expression of COX-2 might lead to the inflammatory cancer phenotype (26, 27) According to several reports, bradykinin contributes to inflammation following SARS-CoV-2 infection (28, 29). Bradykinin levels might be elevated in SARS-CoV-2 patients due to the dysregulation of the angiotensin-converting enzyme levels (29). While it remains to be seen whether blocking bradykinin activity will be useful during SARS-CoV-2 infection, aspirin treatment was reported to be beneficial in SARS-CoV-2 patients (30), warranting a closer examination of the possible effects that other aspirin-based treatments could have on the prevention and treatment of SARS-CoV-2 infection.

Aspirin is already approved as an OTC for sore throat (internal analgesic), yet there are no throat sprays containing aspirin on the market. We have developed a formula containing aspirin and several other food-grade nutritional supplements which act synergistically to help the aspirin act locally and reduce the inflammation and possibly the pain caused by bradykinin. Additionally, we found that when measured against other leading products containing various active ingredients, this formula not only performed better but was much less inflammatory. For one, none of the other active ingredients tested are known to act on bradykinin or its inflammatory pathway. Also, the inactive ingredients in the ASA formula and lozenge (Biovanta) were carefully selected to compliment the anti-inflammatory effects of the active ingredients.

Incidentally, we believe the inflammatory effects of the leading products may have been caused not by the active ingredients but by the inactive ingredients. For example, in Figure 4, four products containing menthol are analyzed, yet the one with the lowest concentration of menthol (Ricola original) shows the greatest increase in PGE2 release and LDH release in the absence of bradykinin stimulation. The products contain numerous inactive ingredients that are difficult to study individually. Inactive ingredients are not closely regulated, and precise concentrations are not usually reported.

The inactive ingredients may have specific or non-specific effects. According to previous reports, hyperosmotic sugar solutions can change respiratory epithelial cell shape and open tight junctions (31). These are the types of morphological effects we show in Figure 6. Also, higher osmolarities can result in the secretion of proinflammatory cytokines (Interleukin-8, Interleukin-6, Interleukin-1β and Tumor Necrosis factor-α) (32), and we did observe increases in IL-8 in response to certain products. We observed significant decreases in TEER, indicating membrane perforation for most of the products tested, which can be caused by osmotic stress. The ASA lozenge (Biovanta lozenge) was formulated with isomalt as the main sugar base, and isomalt has a relatively low osmotic pressure compared to the sugars in the other lozenges tested. To our knowledge, the osmotic pressure of these products on respiratory epithelia has not been measured before.

Taken together, we believe the integrity and inflammatory state of the respiratory membrane epithelium is an important factor to consider in the prevention and treatment of respiratory disease. There are several inflammatory pathways known to be involved and clearly more to be discovered. From our observations, there seem to be several pathways involved in the control of IL-8 expression, including PGE2 and perhaps osmotic stress. We were surprised to learn that many common over the counter products on the market for cold symptoms such as sore throat, greatly exacerbate IL-8 levels. Ongoing studies will determine if this is due to osmotic stress or some other inflammatory signal. In either case, it is highly concerning given the key role IL-8 plays in exaggerating the immune response, thus increasing the likelihood of a “cytokine storm”.

## Acknowledgements

This study was funded by Applied Biological Laboratories.

## Notes

### Competing Interest Statement

This study was funded entirely by Applied Biological Laboratories, a private company that owns the BiovantaTM product. Unless otherwise indicated all experiments were performed at Applied Biological Laboratories research facility located at the SUNY Downstate Biotechnology Incubator, a part of StartUP NY. All of the authors were employees of Applied Biological Laboratories at the time the experiments were performed.

### Summary of Updates

Confirmatory results were added, additional data to support and confirm our findings. Additional data explaining the effects of higher dose aspirin were added.

